# GEMtractor: Extracting Views into Genome-scale Metabolic Models

**DOI:** 10.1101/790725

**Authors:** Martin Scharm, Olaf Wolkenhauer, Mahdi Jalili, Ali Salehzadeh-Yazdi

**Author notes:** Corresponding author: Martin Scharm.

## Abstract

**Summary:** Computational metabolic models typically encode for graphs of species, reactions, and enzymes. Comparing genome-scale models through topological analysis of multipartite graphs is challenging. However, in many practical cases it is not necessary to compare the full networks. The GEMtractor is a web-based tool to trim models encoded in SBML. It can be used to extract subnetworks, for example focusing on reaction- and enzyme-centric views into the model.

**Availability and Implementation:** The GEMtractor is licensed under the terms of GPLv3 and developed at github.com/binfalse/GEMtractor – a public version is available at sbi.uni-rostock.de/gemtractor.

## 1 INTRODUCTION

At the most basic abstraction level, biological phenomena can be represented as mathematical graphs. Genome-scale metabolic models (GEMs) are an example, that describe the associations between genes, proteins, and reactions of an organism. Such models are typically encoded as multipartite graphs, which are partitioned into distinct subsets of vertices representing metabolites, reactions, and enzymes. Since topological analyses of large-scale multipartite graphs is challenging, one is often identifying subnetworks from a whole genome metabolic model. Extracting the reaction-centric (links of reactions) or enzyme-centric (links of enzymes) view simplifies the graph structure and shifts its perspective – from kinetic interactions to phenotypical connections. This opens opportunities for novel topological analyses of the metabolism (Lacroix *et al*., 2008).

The idea of extracting and analysing the enzyme-centric network of a GEM is not new. However, Horne *et al*. (2004) is not available anymore and Asgari *et al*. (2018) apparently mistook the reaction-centric network for an enzyme-centric network.

Here we introduce the GEMtractor, a web-based tool to extract reaction-centric and enzyme-centric views of genome-scale metabolic models (GEM). GEMtractor allows for trimming of models to, for example, remove currency metabolites or to focus on context-based models. The GEMtractor is a free software and easy to install, including an increased privacy and speed because it can be deployed locally.

## 2 TECHNICAL NOTES

The GEMtractor is a Django web application, that we developed with a focus on privacy, compatibility and speed. The interactive front-end is implemented using jQuery and designed using W3CSS. Thus, it works in all modern browsers as well as on mobile devices; there is no need for registration. Figure 1 shows a typical workflow when using the GEMtractor.

**Figure 1.**
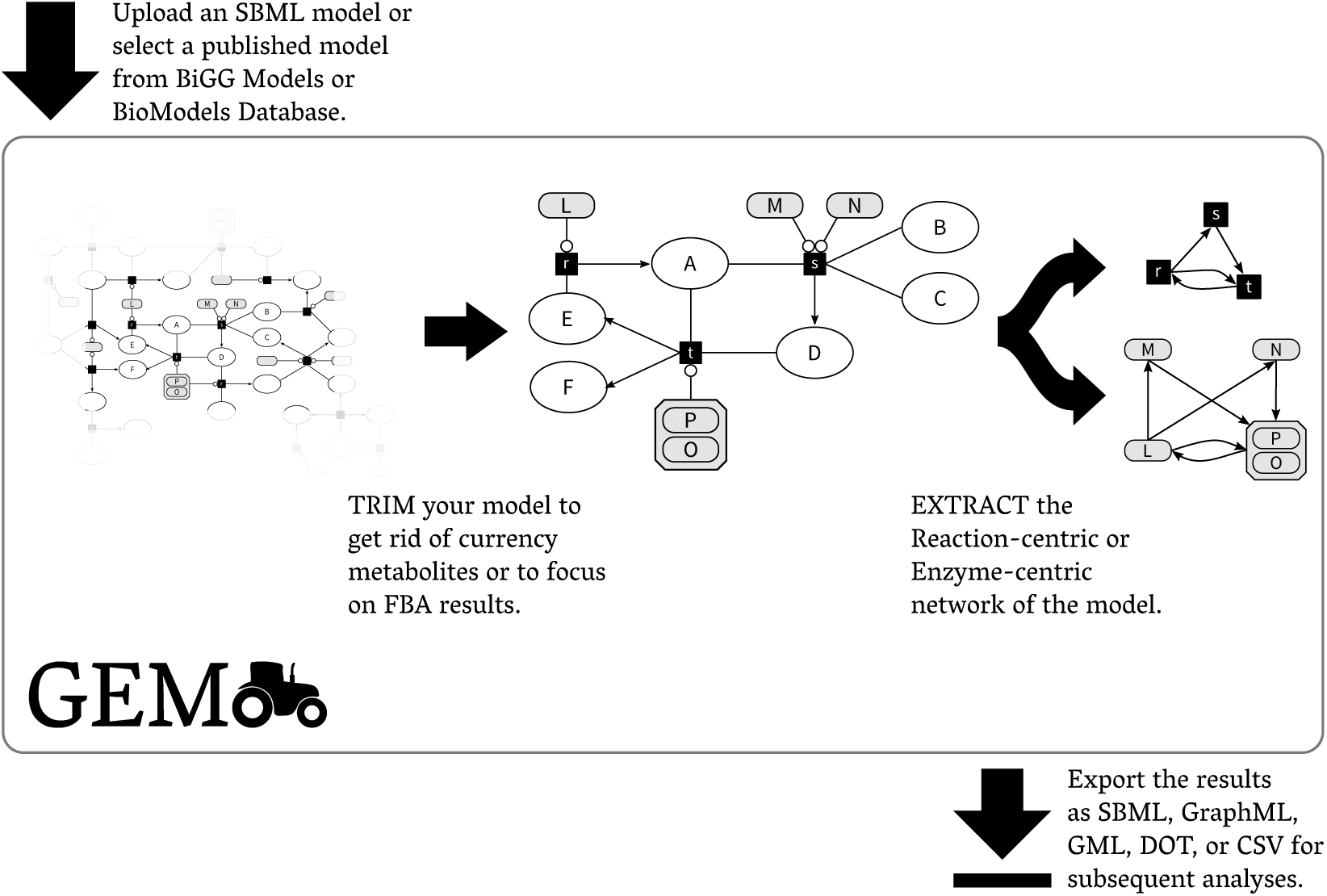
The workflow when using the GEMtractor: A user selects a model, trims undesired entities, extracts a view into the model, and exports the results in exchangeable formats.

The GEMtractor uses libsbml (Bornstein *et al*., 2008) to support all models encoded in SBML. To start an analysis, the user needs to select a model. Users may upload their own models or choose a published model form BiGG (King *et al*., 2016) or BioModels (Chelliah *et al*., 2015). Decoding the metabolite-reaction network from the SBML is straightforward. However, understanding which enzyme combinations catalyse a certain reaction is a bit tricky. The GEMtractor supports two flavours of gene-product associations: Using the reactions’ notes and proper annotations using the SBML FBC package (Olivier *et al*., 2015). Some reactions lack both annotations – in this case, the GEMtractor assumes that there is an implicit enzyme whose identifier matches the identifier of the reaction, prefixed with reaction_–_. We call these invented enzymes *fake enzymes.*

As soon as the model is validly decoded, the user can start trimming entities. The front-end lists all species, reactions, enzymes, and enzyme complexes in individual tables, which can be sorted by different criteria, such as occurences of entity names. It is then comparatively easy to remove currency metabolites. The filters can also programmatically applied using a text-field, so users may save their filters offline or paste filters obtained by their FBA results. Every change on the filters is evaluated at the front-end to check for possible inconsistencies. For example, if an enzyme is to be removed, it may be necessary to also remove all enzyme complexes, in which it participates. Such inconsistencies are highlighted and summarised at the top of the page to inform the user.

The user may then decide to extract a specific view from that trimmed model. In the reaction-centric network consecutive reactions are linked. That means, if a reaction `r` produces a metabolite that is consumed by the reaction `s`, there will be a link from `r` to `s`. Similarly, the enzyme-centric network links consecutive enzymes and enzyme complexes. For example, if `r` is catalysed by `X` *or* `Y` (alternatives, both can catalyse) and s is catalyse by `M` *and* `N` (both are necessary for the catalysis), then there will be two links: From `X` to (`M` *and* `N`) and from `Y` to (`M` *and* `N`). Thus, both the reaction- and the enzyme-centric network are directed, unipartite graphs.

Finally, the obtained graph can be exported into an exchangeable format. The GEMtractor supports SBML, DOT, GML, GraphML, and CSV as output formats. In case of SBML, it preserves as many annotations as possible. For example, the *species* in the reaction-centric network will have all the annotations of the reactions in the original metabolite-reaction network. Similarly, if the gene products were annotated using the FBC package, they will be annotated in the exported file.

In addition to the web browser front-end, the GEMtractor provides a decent API. Trimming and extracting can be encoded in JSON jobs and sent to the API endpoint, which will take care of the computation and immediately return the results. Client implementations in several languages are shipped with the source code. Due to Django’s architecture, the heart of the GEMtractor can be used as a Python module if a network connection is undesired.

Even though the GEMtractor can basically handle all valid SBML models, some models cause problems. For example, the computation time increases with model size, which may cause timeouts at the web server. Similarly the maximum upload size allowed by the web server could terminate a data transfer, so not every model may be upload-able. Furthermore, some models may have invalid gene associations while being valid SBML. That is, because in SBML prior to level 3 there was no standard on encoding the gene associations – they were basically encoded in a free-text field, which is of course error-prone. This is primarily the case for models from BioModels, as those models are often encoded in SBML level 2. If the GEMtractor finds a gene association but is not able to parse it, it will stop and report an error.

While the GEMtractor was built for GEMs, it is applicable to any SBML model. Extracting an enzyme network my be pointless, as kinetic models typically lack the gene association annotations, but you can still trim models to focus on submodels or extract the reaction-centric network for subsequent analyses.

Our public GEMtractor instance at sbi.uni-rostock.de/gemtractor should be useful for the majority of analyses on GEMs. However, if you still need extended quotas, cache options, web server upload limits or timeouts, you are invited to install your own GEMtractor. The installation process is documented on the GEMtractor’s website and in the source code. It requires a Python application server (e.g. gunicorn) to deal with the dynamic pages, and a default web server (e.g. nginx) to serve the static files. However, if you are happy with Docker, installing a GEMtractor boils down to a single command line.

As documentation is key to dissemination, the GEMtractor comes with an extensive FAQ, proper Python documentation and example client implementations in different programming languages. Automatic tests cover most of its source code to prevent future programming mistakes and the GEMtractor exposes a monitoring endpoint to keep an eye on its health.

## ACKNOWLEDGMENTS

We would like to thank Markus Wolfien and Tom Gebhardt for creative brainstormings.

## REFERENCES

V. Lacroix, L. Cottret et al. (2008) An Introduction to Metabolic Networks and Their Structural Analysis, IEEE/ACM Transactions on Computational Biology and Bioinformatics, 5:4, 594–617.

A. B. Horne, T. C. Hodgman et al. (2004) Constructing an enzyme-centric view of metabolism, Bioinformatics, 20:13, 2050–2055.

Asgari Z., Zabihinpour, and A. Masoudi-Nejad (2018) SCAN-Toolbox: Structural COBRA Add-oN (SCAN) for Analysing Large Metabolic Networks, Current Bioinformatics, 13:1, 100–107.

B.J. Bornstein, S.M. Keating et al. (2008) LibSBML: an API library for SBML, Bioinformatics, 24:6, 880–881.

A. King, J. Lu et al. (2016) BiGG Models: A platform for integrating, standardizing and sharing genome-scale models, Nucleic Acids Research, 44:D1, D515–D522.

V. Chelliah, N. Juty et al. (2015) BioModels: ten-year anniversary, Nucleic Acids Research, 43:D1, D542–D548.

B.G. Olivier and F.T. Bergmann (2015) Flux Balance Constraints, Version 2 Release 1, Available from COMBINE identifiers.org/combine.specifications/sbml.level-3.version-1.fbc.version-2.release-1.

